# From tree tops to the ground: reversals to terrestrial habit in *Galeandra* orchids (Epidendroideae: Catasetinae)

**DOI:** 10.1101/224709

**Authors:** Aline C. Martins, Thuane Bochorny, Oscar A. Pérez-Escobar, Guillaume Chomicki, Silvana H. N. Monteiro, Eric de Camargo Smidt

**Affiliations:** Universidade Federal do Paraná, Setor de Ciências Biológicas, Centro Politécnico, Jardim da Américas, Curitiba - PR81531-980. C. P. 19031, Brazil; Universidade Estadual de Campinas, Programa de Pós-Graduação em Biologia Vegetal, Campinas - SP, 13083-970. C.P. 6109, Brazil; Identification and Naming Department, Royal Botanic Gardens Kew, Richmond, TW9 3AB, UK; Department of Plant Sciences, University of Oxford, South Park Road, Oxford OX1 3RB, UK; The Queen’s college, High St, Oxford OX1 4A W, UK; Universidade Estadual de Feira de Santana, Programa de Pós-Graduação em Botânica, Depαrtαmento de Cie□nciαs Biológicas, BR 116Nkm 3, 44031-460 Feira de Santana, BA, Brazil

**Keywords:** Orchids, Amazonia, ancestral area estimation, molecular dating, South American arid diagonal

## Abstract

The colonization of the epiphytic niche of tropical forest canopies played an important role in orchid’s extraordinary diversification in the Neotropics. However, reversals to the terrestrial habit occurred sparsely in species of Epidendroideae. To better understand which factors might have been involved in reversals to terrestrial habits in the predominantly epiphytic Epidendroideae, we investigate *Galeandra* diversification in time and space. We hypothesized that the reversal to the terrestrial habitat is linked to the origin of open vegetation habitats in South America. We generated a time-calibrated phylogeny based on a matrix of 17 out of 20 species of *Galeandra* plus outgroups and seven DNA markers. We found that *Galeandra* originated towards end of the Miocene, about 10 Ma in Amazonia (including the Guiana Shield). The terrestrial clade originated synchronously with the rise of dry vegetation biomes, suggesting that aridification during the last 5 million years dramatically impacted the diversification of epiphytic lineages in the Neotropics. Habit is correlated with floral spur lengths and geographic range size. The longer spurs and narrower ranges are found in epiphytic species: probably adapted to a specialized pollination mode, associated to the long-tongued Euglossini bees, which also prefer forested habits. The terrestrial species presents variable floral spurs and wide distribution ranges, with evidence of self-pollination, suggesting the loss of specialized pollination system and concomitant range expansion.

## Introduction

The colonization of the epiphytic niche in tropical forest canopies played an important role in the extraordinary diversification of orchids in the Neotropics (Benzing, 1987; Gentry and Dodson, 1987; Givnish et al. 2015). Epiphyte microsite specialization together with biotic and abiotic variables including pollinator shifts, CAM photosynthesis, and presence in cordilleras have been proposed as the main drivers of orchid (Givnish et al., 2015; Pérez-Escobar et al., 2017a). The origin of epiphytism in land plants, including orchids, ferns, and leafy liverworts, followed the establishment of angiosperm-dominated canopies in the Cenozoic, possibly facilitated by climate change in the Paleocene/Eocene border (Chomicki et al., 2015a; Feldberg et al., 2014; Schuettpelz and Pryer, 2009)

Orchids are the most diverse group among epiphytes, accounting for 68% (19,000) of the 27,600 vascular epiphyte species (Gentry and Dodson, 1987; Zotz, 2013). The evolution of epiphytism may have enhanced orchid diversification by allowing the conquest of new, largely unoccupied niches (Givnish et al., 2015), and often terrestrial orchid lineages are less diverse than the epiphytic (Gravendeel et al., 2004). The epiphytic habit offers the option of colonizing a larger heterogeneity of habitats: all the surface of branches, twigs and bark. However it requires adaptation to low substrate stability, limited nutrient and water supplies, and to colonize the patchiness biotope (Laube and Zotz, 2003). Thus, the canopy is difficult to colonize and only a plant with a complex suite of adaptations could survive as an epiphyte (Benzing, 1987). Orchid adaptations to tree bark includes root with layer(s) of dead cells known as velamen, which enhance water and nutrient absorption and protects photosynthetic roots against UV-B radiation (Chomicki et al., 2015a), and thick succulent leaves and stems that store water (Freudenstein and Chase, 2015).

In Orchidaceae, epiphytism evolved at least four to seven times over the past 43 million yr and was possibly lost about seven to ten times (Chomicki et al., 2015). In the species-richest subfamily Epidendroideae the epiphytic habit predominates, yet the ancestral condition in orchid is clearly terrestrial (Chomicki et al., 2015; Freudenstein and Chase, 2015). Reversals to the terrestrial habit occurred many times in Epidendroideae, including the Collabieae (Xiang et al., 2014), *Dendrobium*(Xiang et al., 2016), most *Eulophia* (Martos et al., 2014), *Cyanaeorchis* (Batista et al., 2014), *Bletia+Hexalectris+Basiphyllaea* (Sosa et al., 2016), many Pleurothallidinae (Freudenstein and Chase, 2015), Malaxideae (Cameron, 2005) and *Galeandra*(Catasetinae). Although most reversals are associated to species-poor lineages, some are associated to speciose clades, potentially resulting from rapid diversification (Cameron, 2005). The reversals to terrestrial habit are still poorly understood, but they might involve deep changes in morphological adaptations (Zhang et al., 2017) such as the loss or reduction of the velamen (Chomicki et al., 2015a) or occupation of new habitats (Sosa et al., 2016). Some of these adaptations are possibly linked to the presence of AGL12 gene, which are otherwise lost in epiphytic orchids (Zhang et al., 2017).

To better understand the factors that might have influenced the reversal to the terrestrial habits in a linage within the predominantly epiphytic Epidendroideae, we investigate *Galeandra* diversification dynamics, climatic preferences, flower morphology and area of occurrence, using phylogenetic comparative methods. *Galeandra* is a widely distributed genus in the Neotropical region, ranging from southern Florida to northern Argentina, and five of its *ca* 20 species are terrestrial (Monteiro et al., 2010). *Galeandra* species occur across a wide range of biogeographic regions, mainly Amazonia, Cerrado savannahs and Atlantic Forest. Epiphytic *Galeandra* usually occupy more restricted distribution ranges, and occurs in forested areas (e.g. Amazonia), while the terrestrial species occupy open vegetation ecosystems (e.g. Cerrado, excep *G. beyrichii* which inhabits in forest areas, especially altitude forests). The origin of the savannas worldwide, including the south American Cerrado, coincides with the gradual cooling that started at the Miocene climatic optimum (~15Ma) (Zachos et al., 2001), and they fully established about 5 Ma (Simon et al., 2009). Terrestrial *Galeandra* might have occupied these novel environments only recently. This raises the question whether terrestrial *Galeandra* are adapted to cooler and dryer habitats than epiphytic species occurring in rainforests. Further, it is not clear whether the origin of terrestrial *Galeandra*, and potential biome shifts (from forested to open savannas) occurred synchronously to the origin of the Cerrado.

Morphological differences of floral characters can be also observed in terrestrial and epiphytic *Galeandra*. Flowers of epiphytic *Galeandra* possess a long and filiform spur, while terrestrial taxa exhibit flowers with short and saclike spurs (Monteiro et al., 2010). Long floral spurs usually produce nectar or oil, and are accessed by an enlarged part of pollinator’s body (e.g tongues), but also legs (Steiner and Whitehead, 1990; Whittall and Hodges, 2007). Floral spur enlargement is usually linked to pollinators shifts, and it has been hypothesized as an adaptive response to predefined pollinator morphology (Whittall and Hodges, 2007). The extent to which different floral morphologies are associated to distinct habitats is unknown, but habitat preference might have driven the evolution of particular pollination syndromes.

To investigate *Galeandra* diversification in time and space, and potential associated floral trait shifts, we used DNA sequences from Monteiro *et al*. (2010) plus newly generated sequence data of *Galeandra*, including a denser outgroup sampling. We aim to answer the following questions: (*i*) when and where did *Galeandra* originate? (*ii*) When did reversals to terrestrial habits occur, and where and under which habitat? (*iii*) Was habitat shift correlated with shifts in floral morphology, range size or climatic niche?

## Material and methods

### Phylogenetics and dating analysis

Our study builds upon the sampling of Monteiro et al. (2010) for *Galeandra* plus newly generated sequences for *G. leptoceras* and *G. macroplectra*, totalizing 17 (out of 20) species in the genus. The outgroup sampling was also enlarged to better accommodate molecular dating calibrations (see below), comprising representatives of all Catasetinae genera, including *Cyrtopodium* and *Eulophia*, and newly generated sequences of *Catasetum, Clowesia, Cyanaeorchis* and *Cycnoches*. Voucher information and GenBank numbers are presented in Table S1. DNA extraction, PCR conditions and sequencing methods are described in Monteiro *et al*. (2010).

The final matrix consisted of 31 taxa and 6014 nucleotides for five plastid (*ycf1*, psbA-trnH, *rpoB-trnC* and *trnS-trnG*), and two nuclear (xdh, ITS and ETS) markers. Alignments were performed in MAFFT v. 7 (Katoh and Standley, 2013), with default parameters except for the plastid markers which were aligned using the G-INS-i strategy following recommendations for sequences with global homology. ITS and ETS which were aligned using the Q-INS-i strategy, which considers the secondary RNA structure (Katoh and Standley, 2013). The alignments were manually edited in Geneious 6.0 (Biomatters, 2013) to correct obvious alignment errors. In the absence of supported (Maximum Likelihood Bootstrap Support [MLBS] > 75%) phylogenetic incongruence between plastid and nuclear markers, the matrices were concatenated, also in Genious.

Prior to molecular dating analysis, we performed Maximum Likelihood searches and compared our results with a previously published phylogeny of *Galeandra* (Monteiro et al., 2010) and Catasetinae (Pérez-Escobar et al., 2017a). Maximum likelihood tree searches and bootstrapping of the combined dataset using 1000 replicates were performed in RAxML v. 8 (Stamatakis, 2006) using the graphical user interface raxmlGUI 1.3.1 (Silvestro and Michalak, 2012), under the GTR + G model of DNA evolution.

We subsequently time-calibrated our phylogeny, relying on the same matrix of four plastid and three nuclear markers, comprising 31 taxa and 6014 nucleotides, using a Bayesian relaxed-clock approach implemented in BEAST 1.8.3 (Drummond et al., 2012). Absolute divergence times were estimated under the GTR+ G substitution model, and Yule tree speciation model. The Markov Chain Monte Carlo (MCMC) chain was run for 50 million generations, sampling every 10000 generations. Orchids appear to have diverged from the common ancestor of all other members of Asparagales in the Cretaceous around 110 Ma and the crown group ca. 90 Ma, and upper Epidendroids diverged in the Paleogene, around 50 Ma (Chomicki et al., 2015). There are four unambiguous Orchidaceae macrofossils, but none of them is assigned to taxa closely related to Catasetinae (Ramírez et al., 2007). Therefore, the phylogeny of *Galeandra* was secondarily calibrated based on the age obtained by Chomicki *et al*. (2015) for Catasetinae’s crown group of 19.8 (95% Highest Posterior Density Interval [95% HPD]: 14.68-25.73 Ma). These ages are perfectly congruent with (Givnish et al., 2015). A normal distribution prior was applied on the Most Recent Common Ancestor (MRCA) of Catasetinae, with Mean=19.8 and Stdev=3 (95% HPD 14.87 – 24.73). A maximum clade credibility tree was summarized in TreeAnnotator v. 1.8.0 (part of BEAST package) with a 25%burn-in, when effective sample size (ESS) for all parameters were superior to 200, as assessed in Tracer 1.5. Trees were visualized and initially edited in FigTree 1.4.0 (Rambaut, 2009).

### Ancestral area estimation

Species ranges were coded from the literature and from herbarium specimens (ALCB, AMES, AMO, B, BM, BR, F, INPA, K, K-L, NY, P, PORT, RB, S, US and W - abbreviations according to Index Herbariorum (http://sweetgum.nybg.org/science/ih/). Biogeographical areas were derived from literature, as well as from distribution patterns observed in other plant lineages (e.g. Rubiaceae: Antonelli *et al.*, 2009; Bromeliaceae: Givnish *et al.*, 2011; *Cycnoches:* (Pérez-Escobar et al., 2017c). We coded the geographical range of *Galeandra* as: A = Central America, B = Chocó, C = Amazonia, D = Guiana Shield, E = Dry Diagonal and F = Atlantic Forest; Fig. S1 shows coded biogeographical regions. Specimens without reliable geographic locality or with dubious identification were excluded.

For Ancestral Area Estimation (AAE), we relied on the R package BioGeoBEARS (Matzke, 2013), which evaluates several biogeographic models altogether to test for the contribution of evolutionary processes (i.e., range expansion, range extinctions, vicariance, founder-event speciation, within-area speciation) to explain the distribution of modern species. We analyzed independently the models DEC (Ree and Smith, 2008), a modified version of DIVA (Ronquist, 1997), named DIVA-like, and a modified version of BayArea (Landis et al., 2013) or BayArea-like, adding separately the founder-speciation parameter *j*. We assessed the overall fitness of the models conducting likelihood ratio tests based on AICc scores. *Galeandra macroplectra* was excluded from biogeographic analysis due to negative branch lengths.

### Range size

To determine if range size is associated to plant habit, we evaluate distribution ranges of all *Galeandra* species sampled in our phylogeny. Distribution ranges were obtained by literature and herbarium specimens’ examination, as stated in the previous section. We calculated the Extent of Occurrence (EOO) and Area of Occupancy (AOO) for each species using GeoCAT (Bachman et al. 2011, http://geocat.kew.org). EOO represents the defined area contained within the shortest imaginary boundary drawn to encompass all the known sites of occurrence of a taxon, often measured by a minimum convex polygon; AOO represents the area within its “extent of occurrence” which is occupied by a taxon, usually calculated by the sum of all square grids in which the species were registered (IUCN, 2013). Because EOO extrapolates the area of occurrence of a species, we choose AOO for our analysis. The Table S5 shows the values of AOO measured and used in the analysis.

### Ancestral state estimation of spur length

Fifteen out of the 20 know species of *Galeandra* (17 included in our DNA sequence matrix) were included in the analysis. We obtained minimum, mean and maximum values of spur length and width (Figure S1) from herbarium specimens and literature (Monteiro et al., 2010). Whenever possible, we gathered measures from at least five individuals per species. Table S6 provides a list with all measurements of species studied and herbarium specimens sourced. Maximum Likelihood Ancestral State Estimation (ASE) of mean spur length values was conducted using an ultrametric tree derived from dating analysis (see above) and the function *contMap* of the R package ‘phytools’ (Revell, 2012). In addition, to investigated the evolution of spur length through time, we produced a traitgram (Evans et al., 2009) by plotting our ultrametric tree as function of time (from root age to present) and phenotype (i.e. spur length) using the function *phenogram* of the package ‘phytools’. Uncertainties of the ASEs were explored by plotting the probability density of the ancestral estimation in the traitgram.

### Correlation tests

We further test for the correlated evolution between plant habit (0=terrestrial, 1=epiphytic), and spur length under a quantitative genetic threshold model (Wright, 1934; Felsenstein, 2012). This model is applied to discrete variables (e.g. viviparity: Lambert & Wiens, 2013; feeding mode in fishes: Revell, 2013), whose probability of state change is associated to an underlying continuous variable. Correlation analysis was implemented on a Bayesian framework for 1,000,000 generations, with a sampling fraction of 100 generations using the function *threshBayes* in the R package ‘Phytools’ (Revell, 2012).

### Phylomorphospace

To visualize relationship between the spur length, range size and plant habit, while simultaneously accounting for phylogenetic relationship, we generated a morphospace using spur length and range size as continuous variables. To this end, we relied on the function *phylomorphospace* of the R package phytools (Revell, 2012).

### Climatic variables

We mapped 639 georeferenced collection records obtained from floras, GBIF database and herbarium specimens (mean 37, maximum number of record per species 157), and they represent the known distribution of *Galeandra* and extant species included in our taxon sampling. To query GBIF database, we relied on the function *occ* of the R package SPOCC (Chamberlain, 2016). We further extracted corresponding values of elevation and 19 climatic variables (30 seconds resolution) reflecting temperature and precipitation regimes from the WorldClim database (available at: <u>http://www.worldclim.org/current</u>; Hijmans et al., 2005), using the function *extract* of the R package RASTER (Hijmans, 2016).

### Non-metric dimensional scaling analyses

To avoid spurious results arising from inclusion of correlated variable, we determined the Pearson’s correlation coefficients between the bioclim variables and altitude and then included only variables with a Pearson’s correlation coefficient <0.5, taking a single variable in correlated clusters. This way, we selected the bioclimatic variables 1 (Annual mean temperature), 2 (Mean diurnal temperature range), 12 (Annual precipitation), 13 (Precipitation of wettest week), 14 (Precipitation of driest week), and 18 (Precipitation of warmest quarter). We analysed these variables using the R-package VEGAN (Oksanen et al., 2007) to perform non-metric dimensional scaling analyses (NMDS) using the dataset of 657 georeferenced herbarium specimens. To ask whether (i) epiphytic versus terrestrial *Galeandra*, and (ii) short-spurred versus long-spurred *Galeandra* had different niches, we computed the 95% confidence intervals for each group. Overlap between confidence intervals suggests the absence of significant niche differentiation among groups.

## Results

### *Phylogeny* of Galeandra *and time of origin of terrestrial and epiphytic clades*

Our matrix of 31 taxa and 6014 nucleotides for four plastid and three nuclear genes yielded a tree with the same relationships found by Monteiro *et al.*, (2010) analysis for *Galeandra* species (Fig. S1). Our enlarged outgroup sampling scheme represents the genus level relationships in the tribe Catasetinae with high support for the core Catasetinae *sensu* (Dressler, 1983). In our phylogeny, *Galeandra* is sister group to the core Catasetinae (*Catasetum, Clowesia, Cycnoches, Mormodes, Dressleria)* plus *Grobya+Cyanaeorchis* (but larger datasets indicate *Grobya*+*Cynaeorchis* as sister to Galeandra+coreCatasetinae (Batista et al., 2014; Pérez-Escobar et al., 2017a, 2017b, 2016, 2015). *Galeandra devoniana* was recovered as sister to the remaining *Galeandra* species, which is split into a terrestrial and a epiphytic clades, both well supported (ML bootstrap support = 100). Absolute age estimation (Fig. 1, Fig. S2) yielded a root age (i.e. the split between *Eulophia* and Cyrtopodium+Catasetina) of 29 Ma (95% HPD: 17-44). The MRCA of *Galeandra* plus the core Catasetinae age is 17 Ma (95% HPD = 11-23), and the crown group of *Galeandra* originated in the Miocene ca. 9 Ma (95% HPD: 5-13). The terrestrial clade and epiphytic clade diverged from each other in the late Miocene about 7 Ma (95% HPD: 4-11) (Fig 1). The epiphyte clade included species from Amazonia, Mexico, Venezuela, and Guianas, while the terrestrial clade encompassed species occurring in the Dry Diagonal of South America, e.g. Cerrado, and open environments in Colombia and Venezuela. The terrestrial and epiphytic clades crown ages are approximately 4 Ma (95% HPD: 2-6). Most of the adaptive radiation in *Galeandra* occurred in the Pliocene-Pleistocene border (>3 Ma). The Mexican clade of epiphytes diverged 2 (1-5) Ma.

**Fig. 1.**
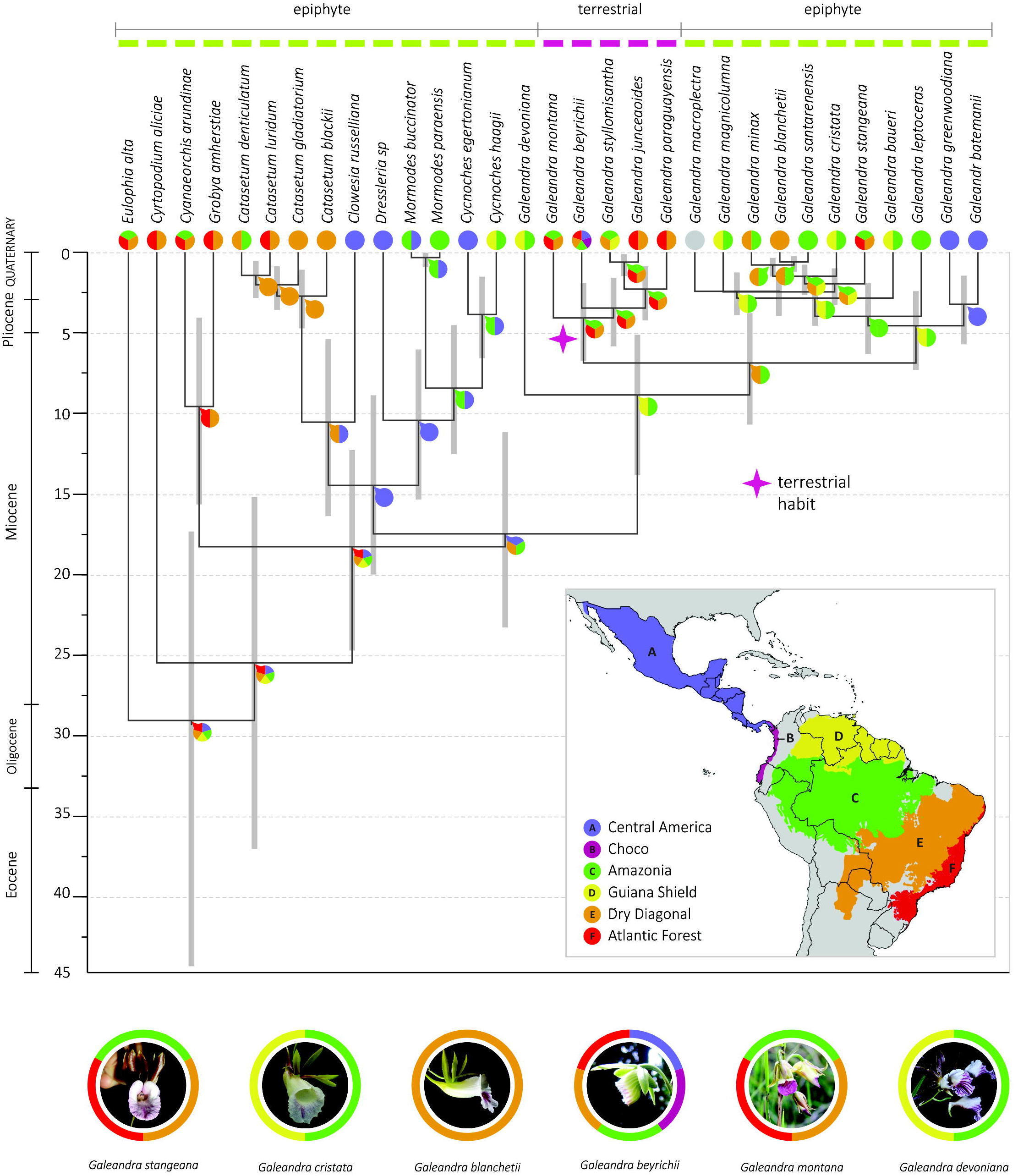
Time-calibrated phylogeny and biogeographic results, showing area coding, and distribution map. Pie charts on nodes represents ancestral area resulted from the BioGeoBEARS analysis and color code follows the legend on the map: A = Central America, B = Choco, C = Amazonia, D = Guiana Shield, E = Dry Diagonal and F = Atlantic Forest. Grey bars represent 95% highest posterior density interval for age estimates. Colored circles on tips represent the occurrence of that species in the delimited geographical areas. *Galeandra macroplectra* presents a grey circle, because it was excluded from the BioGeoBEARS analysis because of negative branch lengths. Photographs in circles represent floral morphological variation in the genus. Colors encircling orchid pictures represent the distribution range of the species according to delimited areas on the map. Photo credits: Adarilda Petini-Benelli (G. *stangeana, G. blanchetii*), Günter Gerlach (G. *devoniana*), Silvana H. N. Monteiro (G. *cristata, G. montana*).

### *Geographic origin of terrestrial habit and biogeographic history of* Galeandra

BioGeoBEARS multi-model approach yielded DEC+j as the best fitting model for the *Galeandra* phylogeny (Tab. S3 provides AICc values of all biogeographical models tested). The MRCA of *Galeandra* is inferred to have lived in an area encompassing Amazonia and Guiana Shield (Fig. 1). The ancestral area for the terrestrial clade, which has at least one widespread species, *G. beyrichii*, included Amazonia, Dry Diagonal and Atlantic Forest. On the other hand, the MRCA of the epiphytic clade is restricted to Northern South America, i.e. Amazonia and Guiana Shield. Among the epiphytic *Galeandra*, only *G. blanchetii* occurs in (and is restricted to) open vegetation of the Dry Diagonal.

### Terrestrial habit origin and correlates with spur length evolution, range size and climatic niche

Figure 2 shows the continuous trait map of floral spur evolution in *Galeandra*,highlighting the marked differences in spur length between the terrestrial and the epiphytic clades. Short spur length was the ancestral condition in *Galeandra* flowers (Fig. 2A), being the shortest in *G. devoniana*. The terrestrial clade shows an intermediate pattern of floral spur length. *Galeandra beyrichi* is the species that presents the shortest spur in this clade. The MRCA of the epiphytic clade had longer floral spurs, and *G. magnicolumna* is the species with the largest floral spurs. The traitgram in Fig. 2B (and density traitgram in Fig. S4) shows the floral spur length mapping in time, highlighting the different rate of evolution of the considered states (short and long spur). We observed a shift in morphological rates at the base of the long-spurred clade of *Galeandra* (epiphytic species) (Fig. 2B). The terrestrial and short-spurred clade presents a higher rate of morphological change.

**Fig. 2.**
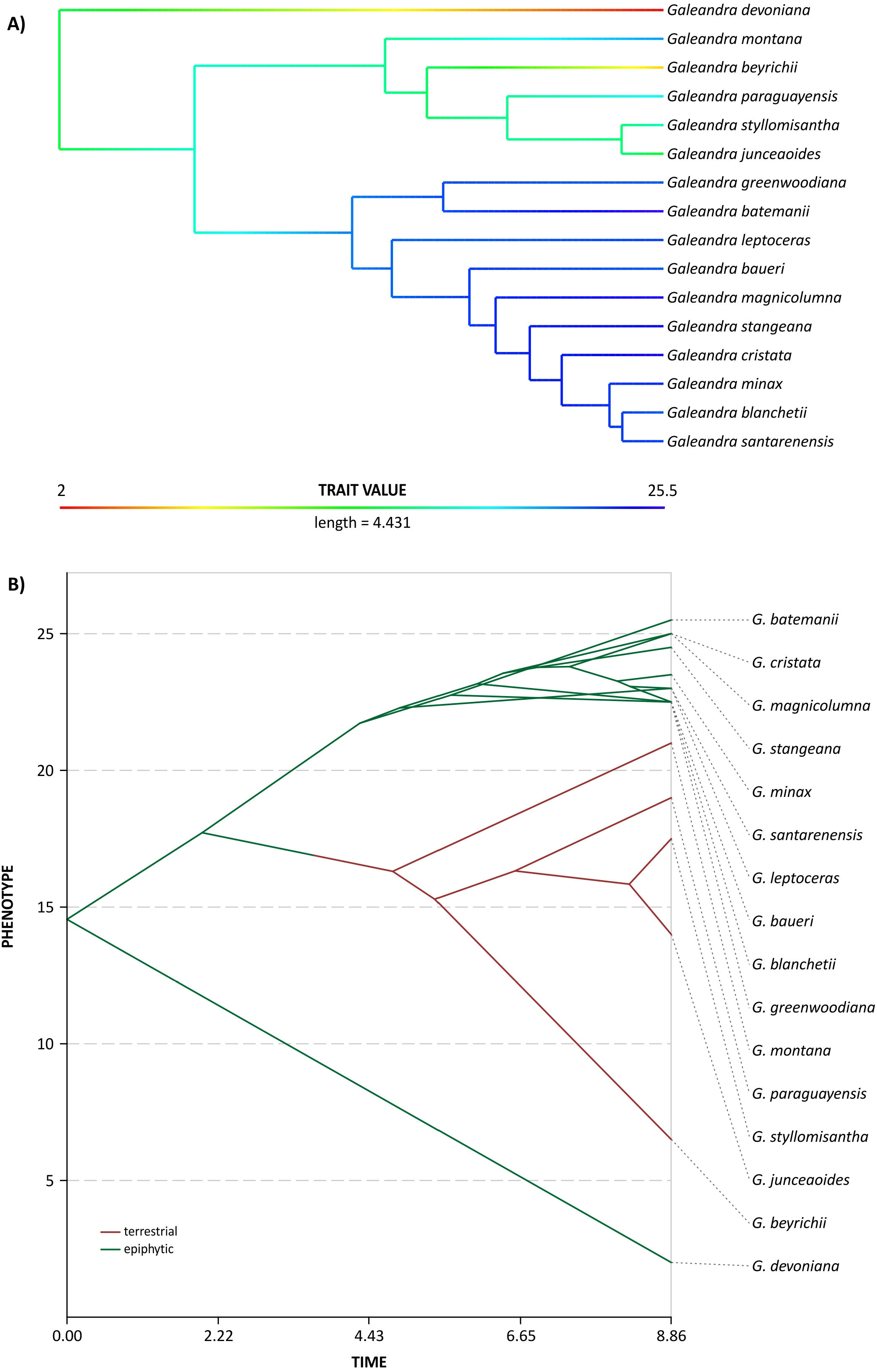
Spur length evolution in *Galeandra* species. A. Ancestral state estimation of spur length in *Galeandra* species; B. Phenogram showing spur length evolution in time.

**Fig. 4.**
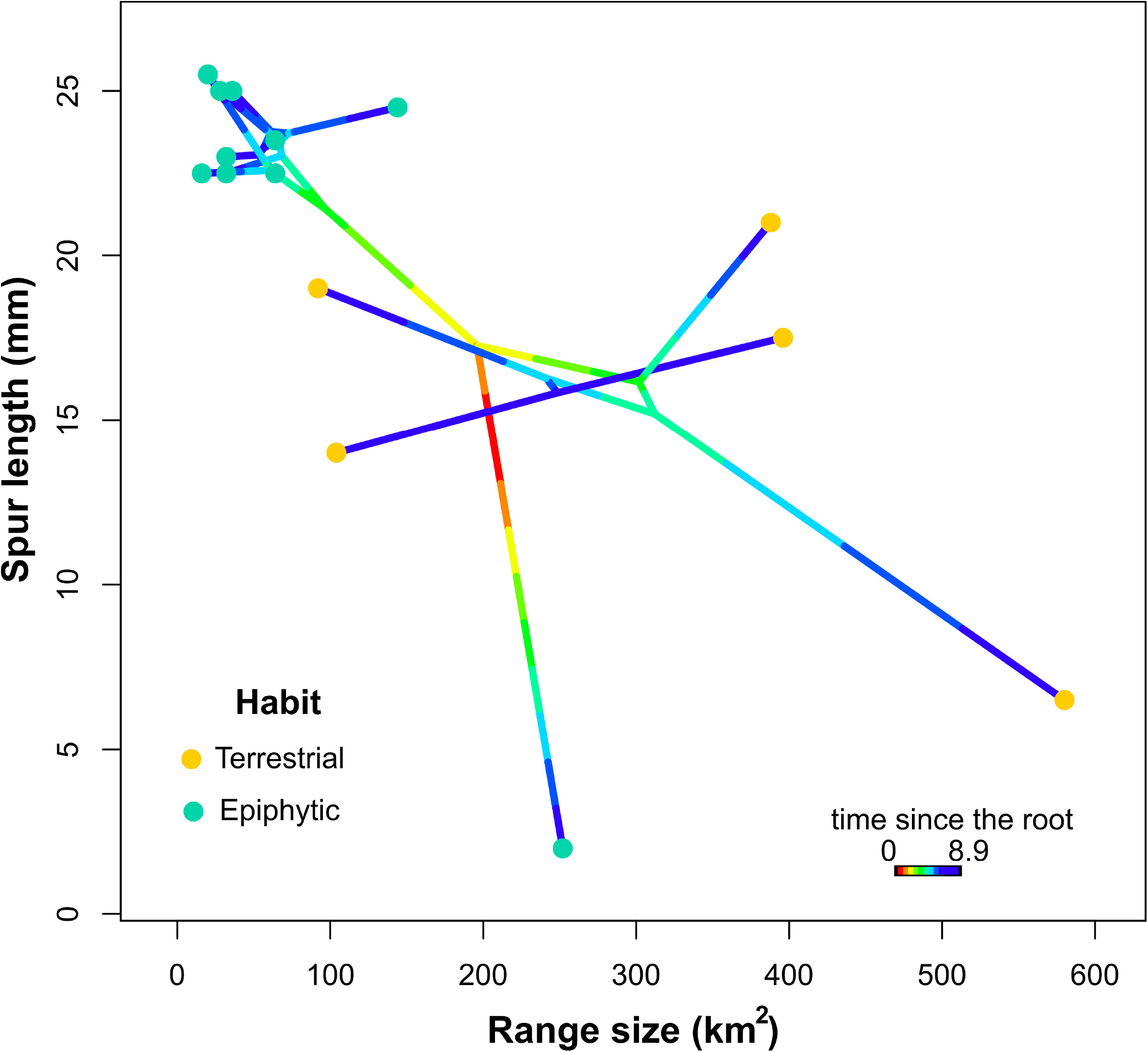
Phylomorphospace analysis result, considering simultaneously spur length and range size as continuous variables, and phylogenetic relationships. Each dot in the graphics represent one *Galeandra* species and the colours distinguish them by habit, terrestrial (yellow) and epiphytic (green). *X* and *Y* axes represent respectively, range size (in km^2^) and floral spur length (in cm).

The figure 3 presents the results of analysis of phylomorphospace correlating floral spur length, range size and habit. Here, epiphytic species formed a cluster of lineages with narrow range sizes (up to 200 km^2^) and longer floral spur (2-2.5 cm). Only one species, *Galeandra devoniana*, does not integrate this group, but presents an isolated pattern of correlation between a short spur length and moderately small range size. The terrestrial species, on the other hand, do not form a cohesive group. They present no pattern of range size or floral spur length.

The NDMS analysis found only the bioclimatic variables 1, 2, 12, 13, 14 and 18 to be non-correlated (Fig. S5). We detected no niche differentiation between terrestrial and epiphytic *Galeandra* neither between long and short-spurred *Galeandra* (Fig. S6 and Fig. S7).

## Discussion

### *Miocene origin of* Galeandra *in Amazonia*

*Galeandra* arose towards the Late Miocene, about 10 Ma, close to the climatic optimum when several plant clades in the Neotropical Region originated (Antonelli et al., 2009; Gustafsson et al., 2010; Hoorn et al., 2010; Pérez-Escobar et al., 2017c). At that time, the Guiana and the Brazilian Shields became large islands. A period of intensified mountain uplift started at the same time in Northern, promoting the origin and extinction of mega wetlands in Amazonia, and shifting the drainage of the Amazon Basin eastwards (Antonelli et al., 2009; Hoorn et al., 2010).

Our ancestral area estimation analyses suggest the MRCA of *Galeandra* inhabited in Amazonia (including the Guiana Shield) towards the Late Miocene, at a period where the region occupied a very large part of South America, extending as far as South Parana region (Hoorn et al., 2010). Early diversification of *Galeandra* occurred in forested areas, while diversification of terrestrial lineages in *Galeandra* might have taken place in both forested areas and the Dry Diagonal, influenced by the rise of dry vegetation biomes. The extend of tropical forests in South America changed with the Andean uplift and its consequences on regional climate, as well as with global climate changes, which favored the establishment of the Dry vegetation areas in South America (Antonelli et al., 2009; Posadas and Ortiz-Jaureguizar, 2016).

The expansion of open vegetation biomes started by the end of Miocene, resulting in the formation of large biomes in the Pliocene such as the Cerrado (Simon et al., 2009), and the establishment of complex system of dry vegetation biomes. Such dry areas are mostly arranged in the dry diagonal of South America, but with accessory patches occurring in all Neotropical Region (Werneck, 2011). In the case of Cerrado, the boundaries appear to have been porous to the migration and recruitment of lineages from a range of wet and dry forest vegetation types. Adjacent biomes including the Amazonia and Atlantic wet forests, tropical and subtropical dry thorn scrub (Caatinga and Chaco), subtropical grasslands, and wetlands have all contributed to the recruitment of Cerrado lineages. The migration of lineages from surrounding ecosystems was facilitated by its nested distribution and enhanced the striking Cerrado’s species richness (Simon et al., 2009). The MRCA of the terrestrial *Galeandra* (4 Ma: HDP 2-6) could have originated on the tropical forests or on the recently formed open vegetation areas. In spite of beginning the diversification about 10 Ma, most plant lineages characteristic of Cerrado diversified only recently, 4 Ma or less (Simon et al., 2009). Therefore, the diversification of both clades (epiphytes and terrestrial) is congruent with this time of drastic transformation in the distribution of forest and open vegetation. However, only species with terrestrial habitat were able to occupy the recently formed biomes, at the same time expanding their distribution. Today, terrestrial *Galeandra* occupy several different open vegetation areas, not only the core Cerrado vegetation, but also costal dunes in Brazil, savannas of Venezuela, islands of savanna in Amazon and even shaded humid forests, in the case of *G. beyrichii*, which probably possess secondary adaptation to forests (Monteiro et al., 2010).

Tropical broadleaf forests provide a plethora of niches for epiphytic plants, and both origins of epiphytism and these forests are profoundly connected (Schuettpelz and Pryer, 2009). Therefore, it is not surprising that the early diversification of the epiphytic clade of *Galeandra* occurred on the forested areas (Amazonia and Guiana Shield), until the Pliocene. In the Quaternary, the Dry Diagonal area became important for this group too. These epiphytes do not occupy the open vegetation, but the gallery forests, which are riverine forests forming narrow strips along the river valleys in the Cerrado biome (Oliveira-Filho and Ratter, 1995). Many plants and animal species from Amazonian or Atlantic Forest domains crosses the Cerrado through those gallery forests, some expanding their distribution within Cerrado (Costa, 2003) and several authors have pointed to affinities between the woody flora of the Cerrado and the Atlantic and Amazon rain forests (Gottsberger and Silberbauer-Gottsberger, 2006; Oliveira-Filho and Fontes, 2000). In the case of *Galeandra*, the gallery forests are occupied by Amazonian species, reinforcing the relatedness of Amazon and gallery forests (Oliveira-Filho and Ratter, 1995; Pennington et al., 2006).

In the epiphytic clade, a dispersal to Central America occurred about 3 Ma (1-5), possibly following the final closure of the Isthmus of Panama, finally formed only 2.8 Ma (O’Dea et al., 2016, but see Jaramillo et al., 2017). The union of both land masses started an exchange of fauna and flora (called GABI or Great American Biotic Interchange – see (Cody et al., 2010)), but for plants there are several evidences of previous dispersals between South and Central/North America (Bacon et al., 2013; Olmstead, 2013). *Galeandra* species might have dispersed overseas as it happened for other orchid species (Renner, 2004), by land, facilitated by the recently formed connection of the Isthmus of Panama or by long distance dispersal across the Andes.

### Habit, floral spur length and range size

Changes from epiphytic to terrestrial habitat might have played a role in *Galeandra* ecological requirements far beyond light and water levels, and may have also affected interactions with bee pollinators. Epiphytic species presents longer floral spurs, which can be associated to the evolution with long tongued pollinators, such as Euglossini bees, the primary pollinators of Catasetinae orchids (Dressler, 1982; Ramírez et al., 2011).

Pollination observations in *Galeandra* flowers are scarce, limited to some punctual observations of pollinaria attached to male orchid bee’s body (i.e. fragrance seeking) (Pearson and Dressler, 1985; Romero-Gonzalez and Warford, 1995) or “Anthophoridae” bees (possibly *Xylocopa*) (nectar or pollen seeking) (Chase and Hills, 1992; Romero-Gonzalez and Warford, 1995). However, how *Galeandra* attract their pollinators remains a mystery. Floral spurs in epiphytic *Galeandra* apparently lack any nutritional reward to pollinators, indicating a possible deceptive attraction, very common in orchids (Ackerman, 1986; Jersáková et al., 2006; Nilsson, 1998; Pansarin and Maciel, 2017). Fragrance-seeking bees can find rewards at least in *G. devoniana, G. magnicolumna* and *G. stangeana* (SHNM pers. obs.) Observations on cultivated epiphytic species of *Galeandra (G. cristata, G. santarenensis, G. stangeana)* shows that they are self-compatible, but not able to self-pollinate, therefore requiring cross-pollination (SHNM pers. obs.).

Terrestrial species have a variable spur length, but in general shorter than epiphytic. The occupation of terrestrial habits might have led to shift in pollinator’s requirements or independence of pollination by animals. Euglossini bees are diverse and widespread in forested habitats, mostly on cloud or lowland forests (Cameron, 2004; Dressler, 1982), presenting low diversity in open vegetation habitats like Cerrado (Faria and Silveira, 2011). The orchid bee fauna occurring in open vegetation biomes are frequently associated to patches of forests occurring along the rivers and there is no species endemic to these biomes, but shared with adjacent large forested biomes like Amazon or Atlantic Forest (Faria and Silveira, 2011). Because flowers of terrestrial *Galeandra* appear to be rewardless, pollination by deceit might also occur in this clade. Also, terrestrial *G. beyrichii* and *G. montana* present very wide distribution ranges (i.e. so and so) and high levels of fruit production in herbaria material, suggesting self-pollination is common among these taxa. However, evidences for self-compatibility, but not spontaneous self-pollination, were observed on cultivated plants of the terrestrial *G. styllomisantha*.

The distribution of orchids might be limited by the joint effect of habitat availability and pollination limitation (Gravendeel et al., 2004; McCormick and Jacquemyn, 2014). Epiphytic *Galeandra* usually presents a narrow geographic range size compared to the terrestrial species. It might be linked to the habit occupied *per se* or to different mechanisms of pollination (dependent of pollinators or not). It could also reflect a higher dependency on particular pollinators. Smaller range sizes are typically found in more specialized mutualisms, as compared to their generalist relatives (Chomicki et al., 2015b). Some evidences of epiphytic orchids with a restricted distribution when compared to the geographic range size of terrestrial species were indicated by Zhang et al. (2015) which related this difference to environmental variables. The low availability of substrates in epiphytic habitats results in restricted and irregular moisture supplies, making water shortages a limiting factor for the establishment and growth of epiphytes (Benzing, 1987; Laube and Zotz, 2003; Zhang et al., 2015). On the other hand, the terrestrial species do not form a cohesive group regarding range size or floral spur length.

Slow morphological change in interaction-related traits is a feature of highly specialized mutualisms, and suggests stabilizing selection (Chomicki and Renner, 2017; Davis et al., 2014). Our morphological analysis shows terrestrial *Galeandra* has a high spur morphorate (Fig. 2), potentially resulting from disruption of bee pollination. As mentioned earlier, long-spurred taxa are likely to have more specialized pollination syndromes than short-spurred ones, which are deemed to be more generalist.

## Conclusions

*Galeandra*, a primarily epiphytic orchid lineage, arose in Miocene about 10 Ma in South America, and most probably the epiphytic clade diversified in Amazon. Terrestrial habit in *Galeandra* arose synchronously with the expansion of open vegetation savannas around 5 Ma. Surprisingly the change from epiphytic to terrestrial habitat does not involve significant changes in climatic niche, perhaps explaining the frequency of such transition in Epidendroid orchids. However, terrestrial species tend to occupy larger geographical ranges probably facilitated by their ecological requirements, but also pollination mode. Floral morphology suggests a shift from pollination by long tongued bees to pollination by morphologically distinct bees or independence from animal pollination.

## Acknowledgements

We are grateful to Adarilda Petini-Benelli, Günter Gerlach, Mauricio Mercadante, Rafael D. Bortoloti and Vinicius Dittrich for *Galeandra* photos. To CAPES for the postdsoctoral fellowship to ACM and CNPq for the Bolsa de Produtividade em Pesquisa (Nível 2 – Proc. 311001/2014-9) awarded to ECS. TB received financial support from CAPES. GC is supported by a Glasstone research fellowship and a Junior Research Fellowship at the Queen’s college, both at the University of Oxford, UK. OAPE is supported by the Lady Sainsbury Orchid Fellowship at Royal Botanic Gardens, Kew.

## Online Supplementary materials

**Fig. S1**. Measurements of *Galeandra* floral spur shown in *G. santarenesis* flower. A. spur width, B. spur length (illustration by A. E. Rocha)

**Fig. S2**. Maximum likelihood phylogeny for the matrix of 31 taxa and 6014 nucleotides for *Galeandra* and outgroups, analysed in RaxML. Values above branches show the ML bootstrap for 1000 replicates.

**Fig. S3**. Time-calibrated tree for the matrix of 31 taxa and 6014 nucleotides for *Galeandraand* outgroups analyzed in BEAST 1.8.3

**Fig. S4**. Traitgram for spur length evolution in time, showing uncertainty about character state using transparent probability density.

**Fig. S5**. Cluster dendogram showing bioclimatic variables relatedness. Bioclimatic variables highlighted in yellow are non-correlated.

**Fig. S6**. Non-metric multidimensional scaling results: epiphytic x terrestrial species.

**Fig. S7**. Non-metric multidimensional scaling results: small x long spurred species.

**Table S1**. List of species used in this study with voucher information and GenBank accession numbers. Newly generated sequences are written in bold. Herbarium acronyms followed the *Index Hebariorum* http://sweetgum.nybg.org/science/ih/

**Table S2**. Botanical collections revised for geographical distribution data of *Galeandra*. Herbarium acronyms followed the *Index Hebariorum* http://sweetgum.nybg.org/science/ih/

**Table S3**. Statistical results from BioGeoBEARS multimodel approach. In bold the best-fit model.

**Table S4**. *Galeandra* species and their Area of Occupancy (AOO) and Extent of Occurrence (EOO), in square meters (km^2^) calculated in GeoCAT (Bachman et al. 2011; http://geocat.kew.org)

**Table S5**. Measurements of floral spur length and width. Herbarium acronyms followed the *Index Hebariorum* http://sweetgum.nybg.org/science/ih/.

